# Rhagovelia uses interfacial run-and-tumble locomotion to improve food capture in flowing environments

**DOI:** 10.1101/2025.04.03.647112

**Authors:** Ishant Tiwari, Nithil Nagappan, Jacob S. Harrison, Saad Bhamla

## Abstract

*Rhagovelia oriander* is a freshwater water strider native to the rivers and streams of North and South America, known for its distinctive skating movement on the water’s surface. This movement resembles the correlated random-walk pattern seen in microorganisms such as *Escherichia coli*. Previous studies have primarily focused on limb adaptations and biomechanics, leaving the ecological significance inadequately addressed. We combine field observations with controlled laboratory experiments using a flow mill to investigate the dynamics of *R. oriander* under typical flow conditions. Our findings indicate that this insect exhibits a two-dimensional run-and-tumble motion, often incorporating lateral tumbles following straight runs (run distance: 30.7 ± 9.3 mm). We find that this behavior is resilient to changes in flow speed. In-silico simulations of particle interception demonstrated that this locomotion method enhances encounter rates compared to linear movement, particularly when the simulated food particle is following a rapid flow field. Our results document run-and-tumble locomotion in a millimeter-scale organism, showcasing a unique example of convergent behavior across diverse taxonomic groups and providing valuable insights into locomotion ecology while serving as a source of inspiration for bioinspired robotics and environmental exploration algorithms.

## Introduction

Navigating complex environments efficiently for exploration and resource acquisition presents key challenges for mobile organisms. Various species from different taxa have developed unique foraging and search strategies (Bell (2012)), often striking a balance between directional persistence and exploratory randomness to enhance their chances of survival and reproduction. Strategies such as correlated random walks (Kareiva and Shigesada (1983)), Lévy flights (Viswanathan et al. (1996); Pyke (2015)), and area-restricted searches (Othmer et al. (1988); Smith (1974)) have been thoroughly studied in both terrestrial and aquatic organisms, showcasing how selective pressures tailor organismal behavior to specific ecological settings.

The run-and-tumble locomotion strategy effectively alternates between straight, directional runs and quick, random reorientations (tumbles). This behavior has been extensively researched in microorganisms such as *Escherichia coli*, which use run-and-tumble to navigate chemical gradients, thus improving their ability to efficiently find nutrients (Berg and Brown (1972); Patteson et al. (2015)). Similarly, the unicellular algae *Chlamydomonas reinhardtii* also employs this strategy by adjusting flagellar synchronization, allowing a shift between straight swimming and rapid reorientations, facilitating effective spatial exploration (Polin et al. (2009)).

Although there has been considerable research at the microbial level (Berg and Brown (1972); Ishihara et al. (1983); Polin et al. (2009); Patteson et al. (2015); Gomez-Marin et al. (2011); Johnson et al. (2024)), run-and-tumble behavior is still largely unexplored at the millimeter scale. *R. oriander*, commonly seen skimming over flowing streams in North and South America, exhibits a movement that resembles the previously described run-and-tumble behavior (Figure 1). *Rhagovelia* is a unique genus within the water strider family (Veliidae). They excel in navigating fast-flowing streams, enabling them to inhabit areas that other water striders do not, and possibly take advantage of new ecological niches (Drake (1914); Cheng and Fernando (1971); Panizzi et al. (2015)). Prior studies have mainly focused on *R. oriander’s* distinct morphological traits, including specialized hydrophilic fans on their middle legs, which enable swift turns through elastocapillary-driven morphing (Cheng and Fernando (1971); Santos et al. (2017); Ortega-Jimenez et al. (2024)). These fans are believed to be the morphological innovation allowing them to thrive in demanding, high-flow environments; however, their unique locomotion behavior might also play a critical role in their success.

**Fig. 1.**
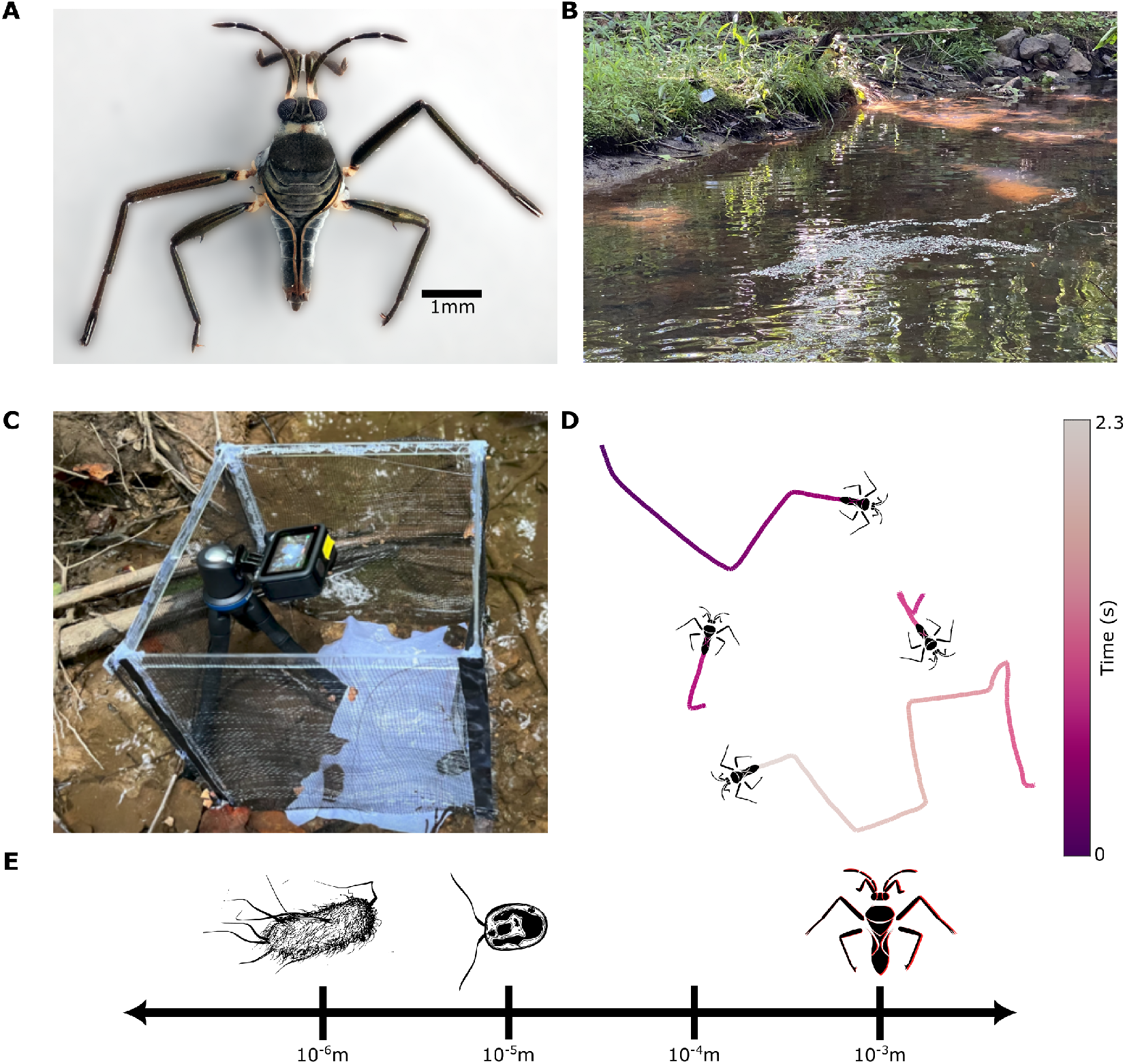
(A) *R. oriander* is a millimeter scale member of the family *Veliidae*. (B) The natural habitat of *R. oriander* consists of shallow streams and creeks, where individuals are typically found near gently flowing water, along edges, and within small eddies or calm surface regions. (C) Environmental observation set up: 10”x10”x10” netted enclosure designed to be dropped over *R. oriander* in its natural habitat (D) Exemplar trajectories of *R. oriander* extracted from field videos show sequential linear motions and sharp turning behavior. (E) Run-and-tumble motion has been observed across a diverse set of taxa and lengthscales in nature, ranging from *E. coli* bacteria and *C. reinhardtii* at the micro-scale to *R. oriander* at the millimeter scale.

In this study, we combine empirical field observations with controlled tests in a custom-designed flow mill to explore how varying flow conditions influence the run-and-tumble behavior of *R. oriander*. We propose that *R. oriander* utilizes run-and-tumble motion to maneuver across the water’s surface, with changes in its runs and tumbles expected to correlate with different flow rates. Furthermore, through mathematical modeling, we anticipate that this behavior will improve the efficiency of encountering potential food particles in simulated flow environments.

## Materials and methods

### Animals

We collected *R. oriander* (Figure 1A) from the East Palisades Trail in Atlanta, GA, USA (33°53’18.8”N 84°26’21.4”W), its natural habitat in ponds and creeks (Figure 1B), and maintained them in a controlled laboratory environment. The insects were kept in a water filled plastic enclosure measuring (12×24×12) inches, and the ambient temperature was maintained at (~20°C). An aquarium filter and air stones were included in the water to ensure a steady flow. The enclosure was supplemented with aquatic plants, algae, and decomposing leaves to ensure that the environmental conditions mimicked their natural habitat. Insects were exposed to a circadian lighting cycle of 12 hours and were fed a diet of live springtails (*Folsomia candida*) daily. Since our study involved working with only invertebrates, it did not require an Institutional Animal Care and Use Committee (IACUC) approval.

### Behavioral Testing

Initial studies were performed at the field site with a GoPro camera mounted on a tripod and a top-down view of a section of the flowing creek (Figure 1C). We selected the recording area by identifying a region with populations of *R. oriander* actively moving on the water’s surface. A subset (n=1-3) of these individuals was enclosed within a mesh boundary that allowed them to remain in the field of view, without obstructing water flow. Characteristic tracks captured during the field recordings are illustrated in Figure 1 D. The movement trajectory of the *R. oriander* displays a unique pattern, characterized by the bug traveling in relatively straight lines, followed by sudden directional changes.

To study the locomotion of *R. oriander* in a controlled laboratory setting, we designed a custom flow mill (Figure 2A) consisting of a three-sided acrylic enclosure (9 x 7 x 2 inches) open at the top. Filtered deionized water was continuously pumped into one side and allowed to flow out over the opposite edge. Below this enclosure, a reservoir housed the pumps for water recirculation and contained an upward-facing light source that was diffused by the acrylic enclosure to create uniform illumination.

**Fig. 2.**
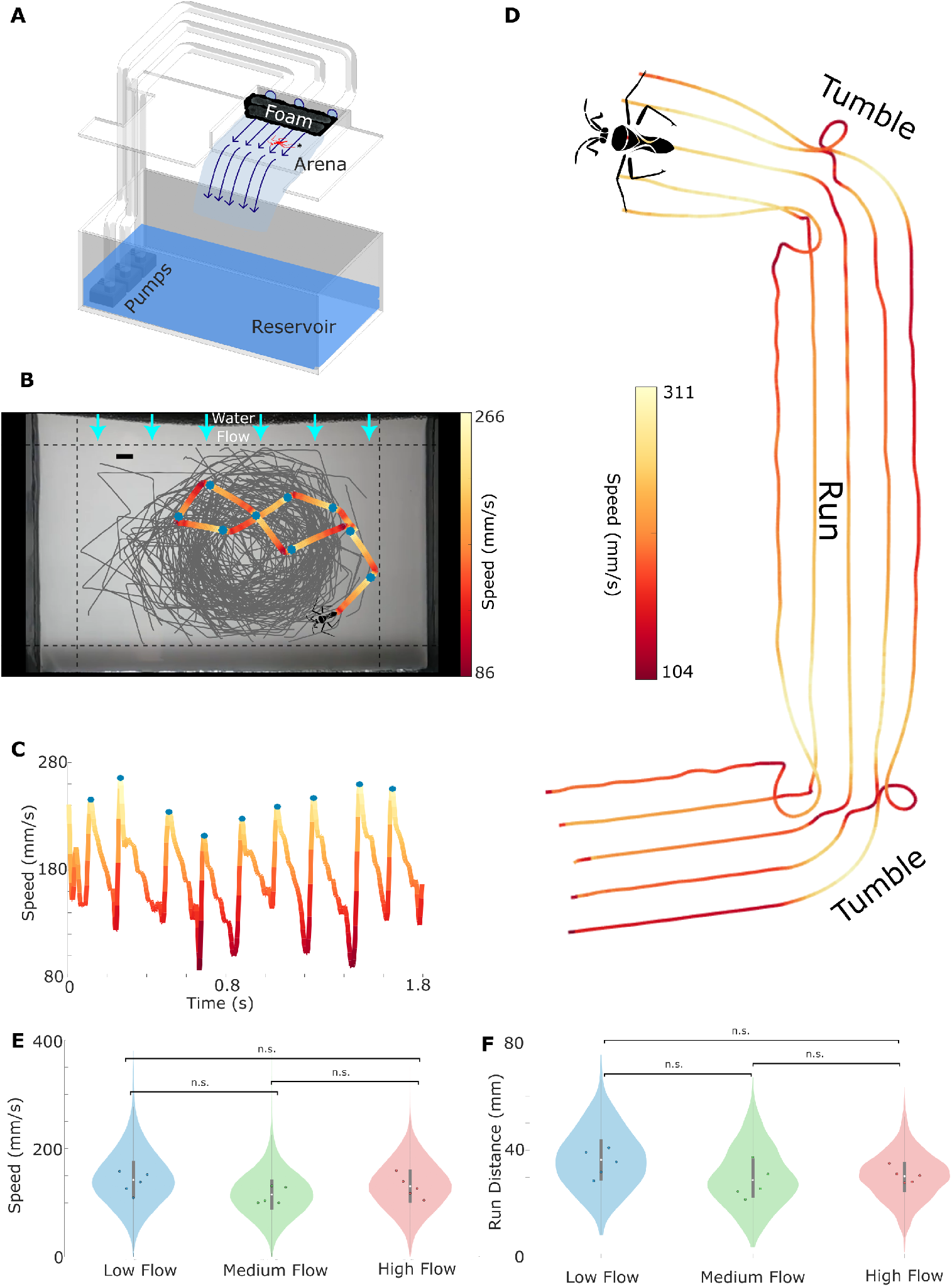
(A) Schematic of the laboratory observational setup of *R. oriander* locomotion. (B) Example snapshot of the experimental arena showing a superposition of insect trajectories. Dashed lines indicate the boundaries used for selecting trajectories included in subsequent analysis. Scale bar, 10 mm. (Insect silhouette not to scale) (See SI video 1 for sample motion of *R. oriander*) (C) Instantaneous speed time series corresponding to the highlighted trajectory in panel B. Blue points indicate local speed peaks, which align with the blue markers on the trajectory shown in panel B. (D) Close-up trajectories of the mid-legs and body center as the *R. oriander* performs a sequence of “tumble–run–tumble” movements. Color gradients along each track represent the instantaneous speed of the corresponding limb. Violin plots showing the distribution of individual (E) median speeds and (F) median run distance across different flow conditions in the arena. Both the quantities were not found to be significantly different across flow conditions (Kruskal-Wallis test for Median speed p>0.1 and Median run distance p>0.09) White dot in the center indicates median of the N=5 replicates. Grey box shows the interquartile range. The whiskers are upper and lower adjacent values.

We verified the velocity fields within the setup using Particle Image Velocimetry (PIV), with lycopodium powder serving as tracer particles to visualize flow trajectories. Flow velocities were adjusted and confirmed at 8–10 mm/s (low flow), 29–31 mm/s (medium flow), and 34–36 mm/s (high flow) before introducing the test individuals to the flow mill. Previous studies (Ortega-Jimenez et al. (2024)) have reported observing *Rhagovelia* in natural flows as high as 1600 mm/s with significant turbulence levels (i.e., ratio between root-mean-square of flow velocity fluctuations to the mean velocity) as high as 65%.

For our experiment, we tested *R. oriander* moving on the surface of the flowing water arena under the three flow regimes (low flow, medium flow, and high flow). Five adult individuals were tested for each flow, resulting in 15 trials, 1 individual per trial. During experimentation, a GoPro Hero 9 was placed viewing the arena top-down and recording the insects at 240 fps, 2K resolution for five minutes.

### Data Analysis

Video recordings of trials were processed using Adobe^©^ Premiere Pro. The video processing included removing fisheye lens distortion and cropping the video to our region of interest. We then used DeepLabCut, a neural network-based markerless tracking software (Nath et al. (2019)), to track two points on the animal: the anterior-most tip of the head and the posterior-most tip of the abdomen, throughout the trial. We extracted the instantaneous coordinates of the head and tail and then took their instantaneous average to determine the approximate location of the body center. From the extracted positional data, we computed various metrics such as instantaneous speed, mean squared displacement curves, run-distance distributions, and turning angles. To reduce noise in our data, we excluded any low-confidence estimates from DeepLabCut as well as any points close to the boundary of the observation arena. The trimmed data yielded periods of trajectory data, which were used for downstream analysis. Exemplary data, along with rejection boundary margins and water flow directions, are visualized in Figure 2B.

### Statistical Analysis

We performed various statistical tests to check for significant differences in distributions across test conditions and across populations. We observed three conditions (low flow, medium flow, and high flow conditions) with n=5 individual replicates for each flow.

For linearly distributed metrics such as speed, run distance, and mean squared displacement (MSD) exponents, we conducted the following statistical analyses. We quantified the variability within each flow condition by computing the standard deviation (SD) of the time series for each individual replicate and then applied Levene’s test (Levene (1960)) to evaluate differences in variance across flow conditions. We then computed the per-replicate medians and conducted non-parametric Kruskal–Wallis tests (Kruskal and Wallis (1952)) to assess whether the median values of these metrics differed between conditions. If a significant main effect emerged, we performed post hoc pairwise comparisons using the Bonferroni correction to account for multiple testing. To directly compare MSD scaling exponents at short versus long timescales within individuals, we employed the Wilcoxon signed-rank test (Wilcoxon (1992)), a non-parametric paired test appropriate for small sample sizes and non-normally distributed data. In addition to reporting the *p*-values, we quantified effect sizes using Cliff’s delta (Cliff (1993)) (*δ*), providing a robust, non-parametric measure of the magnitude of observed differences between short- and long-time MSD exponents. In addition, we evaluated whether the step-length distributions followed a scale-free (power-law) form by applying the Clauset–Shalizi–Newman goodness-of-fit procedure (Clauset et al. (2009)), which estimates the tail exponent, and computes a Monte-Carlo *p*-value. Subsequently, to characterize the underlying distributional shape when power-law fits were rejected, we compared alternative light-tailed models (Weibull, Gamma, and Lognormal) using maximum likelihood estimation and model selection based on the Akaike Information Criterion (AIC) (Akaike (2003); Anderson and Burnham (2002)).

We applied the Hermans–Rasson (Hermans and Rasson (1985)) test for each condition on the pooled heading data to assess directional uniformity, which is sensitive to multimodal deviations and well-suited for detecting non-uniform heading distributions. Circular statistics were calculated using the CircStat toolbox (Berens (2009)). Circular mean, variance, and skewness were calculated for each replicate in a flow condition, and the differences of these statistics across flow conditions were estimated using the Kruskal-Wallis test. Post hoc tests were only performed if there was a significant difference. Together, these analyses allowed us to evaluate both linear and directional behavioral metrics as a function of flow condition, accounting for variability within and across individuals.

### Simulations

We simulate the insects’ motion using a correlated random walk model. The simulations were performed using MATLAB programming language. The complete code for this simulation is available on GitHub (Tiwari et al. (2025)).

## Results

### Kinematics

We observe that *R. oriander* generates bursts of forward motion by actively rowing with its middle legs. Each stroke produces a sharp increase in speed (from ~100mm/s to ~300mm/s), marking the start of a propulsion phase. Immediately after, the insect enters a passive coasting phase during which it decelerates smoothly while maintaining a relatively straight trajectory. Following each run, *R. oriander* typically executes a sudden directional change or tumble. These tumble events occur at the end of the deceleration experienced during the insect’s linear motion. The insect reorients its body, often by several tens of degrees during a tumble, in what appears to be a random direction. It then swiftly resumes motion by rowing with its mid legs to initiate a new run in the updated direction. This rapid (~5 rows per second), recurring pattern of propulsion followed by glide creates a characteristic speed-time series: sharp peaks corresponding to rowing events followed by gradual declines in speed. The profile presented in Figure 2C is consistent across individuals and flow conditions, reflecting the biomechanical rhythm underlying this insect’s locomotion. Previous work (Ortega-Jimenez et al. (2024)) have also reported that these insects are actively swimming in this fashion for most of their lifetimes. Figure 2D depicts a close-up visualization of this run-and-tumble behavior by overlaying the trajectory of the insect’s center body along with the midleg segments, specifically the middle joints and terminal tarsi. The trails are color-coded by their respective instantaneous speed, revealing that peak body velocity occurs shortly after the tumble, as the insect initiates propulsion in the new direction. The figure indicates two distinct tumbles and one run, highlighting the sharp reorientation between runs and the biomechanical coordination involved in each transition. This visualization confirms that run-and-tumble behavior in *R. oriander* emerges from coordinated leg dynamics that alternate between propulsion and directional change.

### *R. oriander* locomotion is independent of tested flow rates

To assess whether external fluid forces affect locomotor output, we quantified the distribution of insect speeds and run distances across three flow regimes. Figure 2E shows the distribution of median speeds for individual insects under low, medium, and high flow conditions. The violin plots reveal overlapping distributions, and statistical analysis using the Kruskal-Wallis test indicates that there is no significant difference between conditions (*p >* 0.1).

We similarly compared the distances covered during each run phase across flow conditions. As shown in Figure 2F, the distribution of run distances remains consistent, with a typical run length of approximately 30 mm across all tested flows. Once again, no significant differences were observed (*p >* 0.09), suggesting that *R. oriander* maintains its locomotor pattern regardless of ambient flow speed. *R. oriander* ‘s run-length distributions were also assessed for scale-free properties indicative of Lévy walks. However, power-law fits were rejected in all individuals across conditions (Clauset–Shalizi–Newman test, *p <* 0.001), and step-length distributions were instead better described by light-tailed models, such as the Weibull or Gamma distributions (See SI for model comparison using AIC). This confirms that the insects’ run-and-tumble trajectories are not Lévy-like, but rather follow a bounded statistical structure that remains robust across flow conditions. These findings indicate that the insect’s run-and-tumble strategy is not only stable across the tested flow environments but also governed by finite-length excursions rather than scale-free locomotion.

### *R. oriander* performs Run and Tumble locomotion

To quantitatively characterize the movement of *R. oriander*, we calculate the mean squared displacement (MSD) as a function of time (Figure 3A). On a log-log scale, the MSD initially rises approximately *t*^2^ at short timescales (*t <* 0.25 s), indicating ballistic motion. However, beyond 0.25 seconds, the MSD growth slows, transitioning into a diffusive/sub-diffusive regime, marked by exponents around 1. This shift represents a qualitative reduction in effective movement speed over longer durations. Importantly, the observed timescale for this transition closely corresponds to the average interval between consecutive rowing events (approximately 0.25 seconds).

**Fig. 3.**
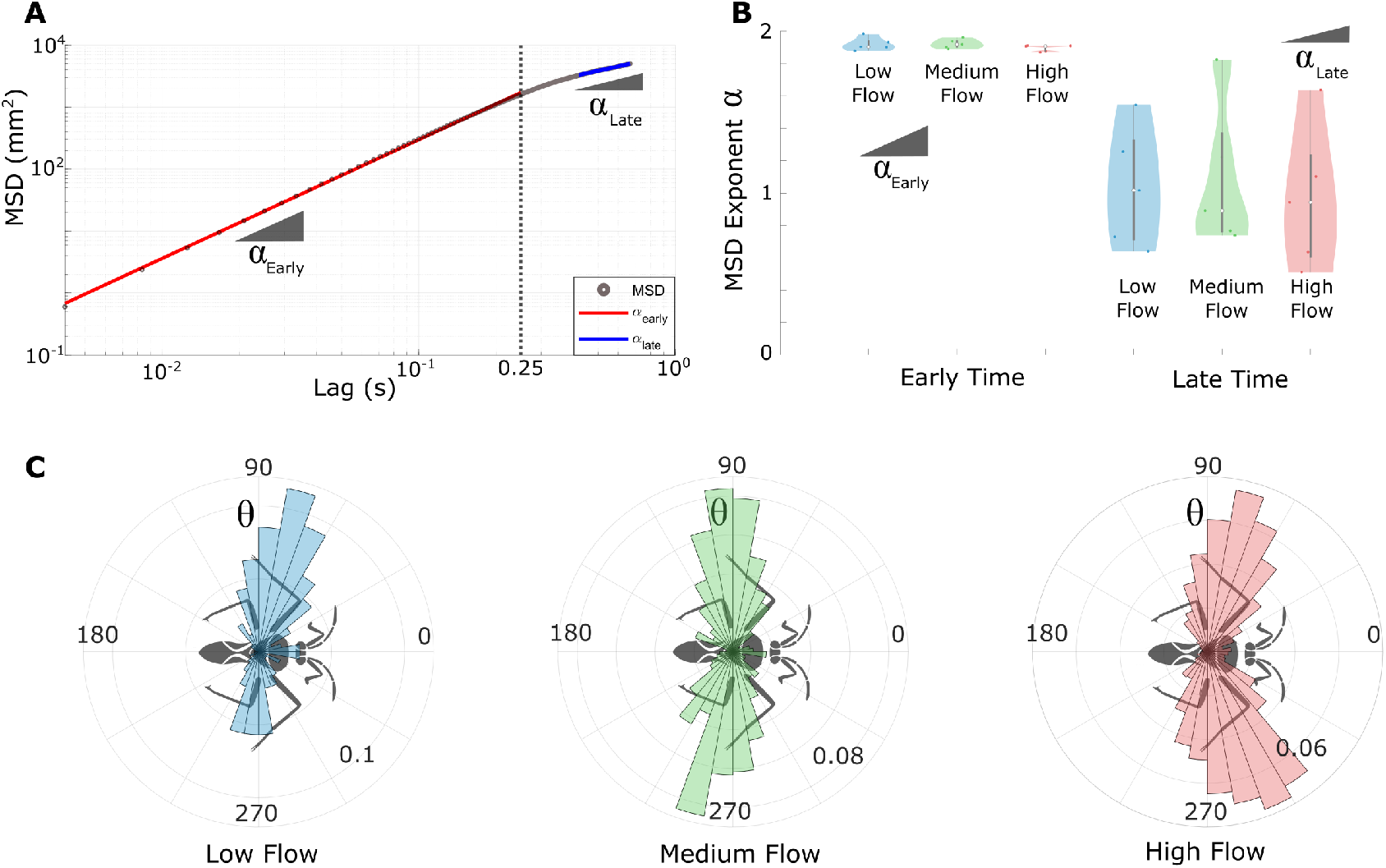
(A) Mean squared displacement (MSD) as a function of lag time shows an early-time scaling exponent of approximately 2, indicative of ballistic motion, followed by a reduced exponent around 1 at later times, consistent with diffusive/sub-diffusive behavior. (B) Early-time exponents (*α*_Early_) are significantly higher than late-time exponents (*α*_Late_) across all flow conditions, indicating a shift in locomotor behavior at longer timescales. (Wilcoxon signed-rank tests; low flow: *p* = 0.063, Cliff’s *δ* = 1; medium flow: *p* = 0.063, Cliff’s *δ* = 1.00; high flow: *p* = 0.063, Cliff’s *δ* = 1.00). White dot in the center indicates median of the N=5 replicates. Grey box shows the interquartile range. The whiskers are upper and lower adjacent values. (C) Polar histograms of *Rhagovelia oriander* ‘s turning angles show a strong preference for lateral left and right turns. (Herman-Rasson test for multimodality in circular distributions; low flow: *p <* 0.0001, *T* = 2.94 × 10^5^; medium flow: *p <* 0.0001, *T* = 1.04 × 10^6^; high flow: *p <* 0.0001, *T* = 2.69 × 10^7^) The angular distribution is consistent across all flow conditions when the N=5 replicate means of circular statistics were compared.(Kruskal-Wallis test applied: on mean vector length: *p* = 0.1; on circular variance: *p* = 0.1; on circular skewness: *p* = 0.23)

To further validate this behavioral shift, we compare the MSD scaling exponents calculated at short and long timescales across all three flow conditions (Figure 3B). In each flow regime (low, medium, and high), the average MSD exponent consistently decreases from short to long timescales, indicating a steady transition from ballistic to diffusive/sub-diffusive behavior. Wilcoxon signed-rank tests showed a consistent downward shift in the MSD exponent across all flow conditions, trending toward statistical significance in each case (low flow: *p* = 0.063, Cliff’s *δ* = 1.00; medium flow: *p* = 0.063, Cliff’s *δ* = 1.00; high flow: *p* = 0.063, Cliff’s *δ* = 1.00). Considering that the typical run distance for the *Rhagovelia* in Figure 2F is approximately 30 mm, while the observation arena lengthscale is almost 6 times the typical run-length, we can safely assume that the decrease in MSD slope is not appearing due to boundary constraints. This observed decrease in slope can also appear if the insect drastically switches direction after each run, thus plateauing the MSD growth after only a relatively short time of a single run. This evidence quantitatively confirms the dynamic locomotion patterns of the insect characterized by alternating run-and-tumble movement. The MSD crossover alone is insufficient to support a Lévy-flight interpretation, as similar ballistic-to-diffusive transitions also occur in correlated random walks. However, when combined with the step-length analysis (Figure 2F), which reveals light-tailed distributions (Weibull or Gamma) and rejects power-law fits across all individuals (*p <* 0.001), the evidence strongly supports a correlated random walk rather than a Lévy flight. Thus, while the short-time scaling is consistent with persistent (ballistic) motion, the overall pattern is finite, and not scale-free.

To confirm that the observed correlated random walk involves sharp reorientations characteristic of run-and-tumble behavior, we next analyze the turning angles between consecutive runs (Figure 3C). A polar histogram (rose plot) of turning angles reveals a strong preference for sharp directional reorientations, with approximately equal probabilities of turning left or right. (See SI for speed dependence of the *Rhagovelia* as a function of direction of its motion with respect to the water flow.) To quantitatively test this multimodal distribution, we apply the Hermans–Rasson test for circular uniformity. The test strongly rejects uniformity for pooled turning angle data under all flow conditions (low flow: *p <* 0.0001, *T* = 2.94 × 10^5^; medium flow: *p <* 0.0001, *T* = 1.04 × 10; high flow: *p <* 0.0001, *T* = 2.69 × 10), confirming a significant tendency for lateral directional changes. Furthermore, we test whether the turning angle distributions vary across flow regimes by calculating circular statistics (mean vector length *R*, circular variance, and circular skewness) per insect and applying the Kruskal–Wallis test. No significant differences emerge among flow conditions (mean vector length: *p* = 0.1; circular variance: *p* = 0.1; circular skewness: *p* = 0.23). Together, these analyses strengthen the characterization of *R. oriander* ‘s locomotion as consistent run-and-tumble behavior, defined by periodic runs punctuated by distinct lateral reorientations.

### Run-and-tumble enhances interception of drifting particles

Having established that *R. oriander* employs a run-and-tumble locomotion strategy, we now investigate the potential ecological advantages of this behavior. To address this, we develop a numerical simulation to examine how run-and-tumble dynamics affect encounter rates with passively drifting particles, which act as inert proxies for potential food items.

The simulated *R. oriander* moves at speeds drawn from a Gaussian distribution based on experimental observations and executes runs of approximately uniform duration. After each run, the simulated insect experiences directional changes modeled by a superposition of two circular Gaussian distributions with means separated by an adjustable parameter Δ*ϕ* (Figure 4A,B). Specifically, a value of Δ*ϕ* = 0° corresponds to predominantly straight movement, while Δ*ϕ* = 180° indicates frequent sharp turns that are equally likely to be left or right, with some finite variance around ±90°. Characteristic turning angle distributions and representative trajectories for Δ*ϕ* = 0°, 90°, and 180° are shown in Figure 4A,B.

**Fig. 4.**
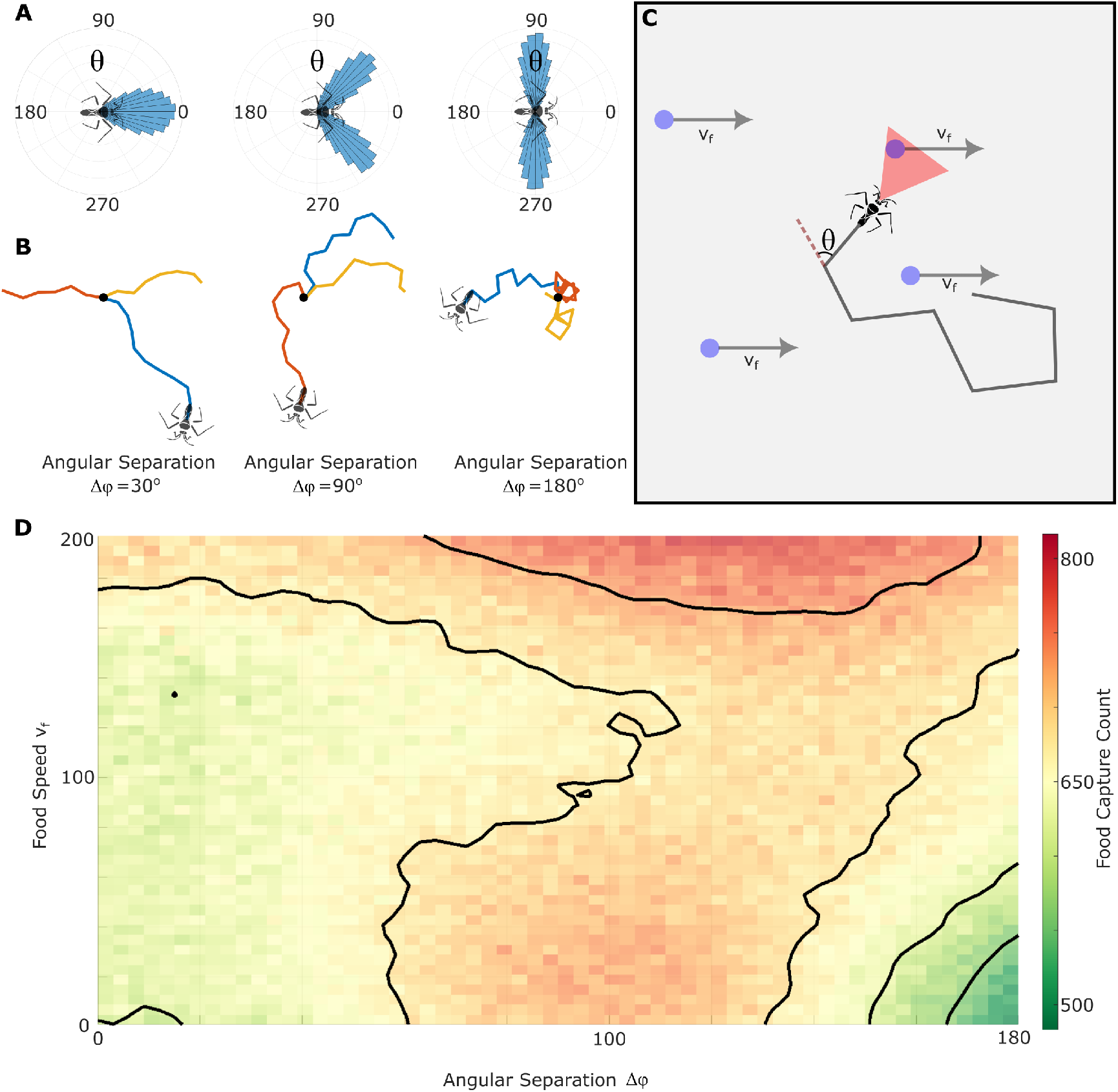
(A, B) Sample turning angle distributions and simulated trajectories for Δ*ϕ* = 0°, 90°, and 180°, illustrating increasing angular reorientations. (C) Schematic of the simulation setup showing the vision triangle of the predator and food particle drifting from left to right. (D) Heatmap of particle encounter count as a function of Angular Separation (Δ*ϕ*) and particle speed (*v*_*f*_), averaged over 10 simulations. Particle encounter is maximized at intermediate Δ*ϕ* values for slow particles, while faster food particles favor sharper directional changes (Δ*ϕ* ≈ 130°–150°), indicating an advantage of run-and-tumble-like strategies in dynamic environments.

In the simulation environment, particles passively drift from left to right within a square domain of side length *L* under periodic boundary conditions, moving at a constant horizontal speed *v*_*f*_ (Figure 4C). This particle speed *v*_*f*_ serves as a second adjustable parameter in our model. The simulated predator is equipped with a forward-directed “vision triangle,” defined as an isosceles triangle with a vertex angle of 45 degrees and side lengths of 40 mm, reflecting the typical run length observed experimentally. Particles are considered intercepted when they enter this triangular visual region.

We explore the effects of turning sharpness Δ*ϕ* (0–180 degrees) and particle speed *v*_*f*_ (0–200 mm/s) on encounter success, quantifying the number of particles captured over fixed-duration simulations. Each parameter combination is evaluated across ten replicate simulations to ensure robust statistical characterization of model performance.

We visualize the combined effect of turning sharpness (Δ*ϕ*) and particle speed (*v*_*f*_) on encounter success using a two-dimensional heatmap (Figure 4D). The x-axis represents angular separation Δ*ϕ*, the y-axis corresponds to particle speed *v*_*f*_, and the color encodes the total encounters during the simulations. At low particle speeds, capture rates show a clear unimodal relationship with Δ*ϕ*, peaking around Δ*ϕ* ≈ 100°. Notably, at these low speeds, extreme directional changes (Δ*ϕ* ≈ 180°) result in the lowest particle encounter rates, performing even worse than straight-line movements (Δ*ϕ* ≈ 0°). As particle speed increases, however, this peak in particle encounter shifts toward higher angular separations (Δ*ϕ* ≈ 130°–150°), suggesting that more pronounced lateral turning becomes increasingly advantageous as food particles move faster.

These results indicate that while moderate lateral turning optimizes number of encounters when particles move slowly or remain nearly stationary, sharper directional changes become increasingly beneficial as particle speed increases. Therefore, the run-and-tumble strategy employed by *R. oriander*, characterized by frequent lateral turns, likely represents an adaptive response that enhances food particle encounters in environments with flowing substrates.

## Discussion

In this study, we document and quantitatively characterize the run-and-tumble locomotion exhibited by *R. oriander*, an interface-dwelling insect navigating dynamic, fluid environments Our observations demonstrate that *R. oriander* employs repeated cycles of rapid directional propulsion (runs) interspersed with sharp lateral reorientations (tumbles), a behavioral pattern well-documented in microbial taxa such as *E. coli* and *C. reinhardtii*, but seldom observed at macroscopic scales. Through controlled laboratory experiments and computational simulations, we reveal that this distinctive locomotion pattern likely provides ecological advantages by augmenting prey encounters, particularly when prey are found on a moving or flowing substrate.

The behavior of *R. oriander* is a compelling example of behavioral convergence. Despite significant morphological and ecological differences, diverse taxa ranging from microscopic bacteria and algae to interfacial insects at millimeter scales independently evolve similar locomotion strategies to efficiently search for resources or more favorable environments. The persistence of run-and-tumble movement across a wide range of body sizes underscores its relevance as a search strategy, aligning with expectations from scaling principles, particularly in environments where continuous linear motion may be less effective.

Run-and-tumble locomotion in *R. oriander* persisted across ambient flow conditions found in its natural environment. Our research shows that essential locomotion parameters, including speed and run length, remain largely unchanged across the ecologically relevant flow regimes we explored. This stability likely allows *R. oriander* to take advantage of different microhabitats near gentle stream edges and within small eddies or surface areas, areas where prey availability and fluid dynamics can vary unpredictably. Such resilience in behavioral strategies is important for organisms living in fluid environments marked by varying external conditions.

From a broader ecological perspective, the run-and-tumble strategy observed here serves as a fascinating evolutionary solution for effective spatial exploration and resource gathering. Previous theoretical and empirical research has recognized search patterns such as Lévy flights and correlated random walks as advantageous in specific scenarios. Our research contributes to this ongoing conversation by showcasing run-and-tumble locomotion in the aquatic interfacial environment, which is particularly suited to the dynamic and complex conditions faced by mobile organisms.

Beyond fundamental ecological insights, the behavioral mechanisms of run-and-tumble locomotion documented in *R. oriander* hold significant potential for biomimetic technological applications. Specifically, the observed locomotion patterns can inspire the design of search algorithms for autonomous surface robots. Such biomimetic algorithms could be particularly valuable in developing mobile swarms (Rubenstein et al. (2014)) for applications including oceanic plastic debris capture, oil spill mitigation, and search-and-rescue operations (Murphy (2007)). Robots operating at fluid interfaces must efficiently explore large spatial areas while adapting to complex surface flows and unpredictable targets, conditions analogous to those encountered by interfacial insects. Therefore, embedding principles derived from *R. oriander* ‘s run-and-tumble locomotion into algorithmic control frameworks could lead to enhanced operational efficiency, robustness, and adaptability of autonomous robotic systems.

In conclusion, we recognize several limitations within our study and list some future avenues. Although laboratory experiments are systematic and controlled, they might not completely reflect the intricacies of natural habitats. Moreover, our computational model streamlines prey dynamics and predator sensory systems, presenting room to integrate more realistic ecological interactions and environmental variability in future simulations. For example, we could incorporate escape tendencies and active motion in the simulated prey, along with a feedback-controlled driver for the predator to actively adapt its run and tumble behavior based on its sensory inputs. Given that *Rhagovelia* have been observed preying upon microcrustaceans and small insects (Drake (1914); Panizzi et al. (2015)), which exhibit varying levels of active locomotion in addition to passive drift, future research could focus on characterizing the movement trajectories and behavior of these prey items. Investigating how *Rhagovelia* detect and actively pursue such mobile prey in flowing water would provide deeper insights into their foraging strategies and ecological interactions. Pursuing these proposed extensions will also expand possibilities for bioinspired robotics and adaptive computational algorithms.

## Supplementary data

Supplementary data available at *ICB* online.

## Author contributions statement

I.T., J.S.H. and S.B. conceptualized the research. I.T., N.N. and J.S.H. designed the experiments. N.N. conducted the experiments, for which I.T. performed the analysis. I.T. did the simulations. S.B. supervised the research. All authors contributed to writing, discussion, and revising the manuscript.

## Acknowledgments

S.B. acknowledges funding support from NIH Grant R35GM142588; NSF Grants MCB-1817334; CMMI-2218382; CAREER IOS-1941933; PHY-2310691 and the Open Philanthropy Project. Text in this paper was revised using LLMs.

