## Supplementary material for "Rhagovelia uses interfacial run-and-tumble locomotion to improve food capture in flowing environments": SI text

*Integrative and Comparative Biology*, 2023, pp. 1–1

doi: DOI HERE
Advance Access Publication Date: Day Month Year
Symposium

SYMPOSIUM

### Supplementary Information: Rhagovelia uses interfacial run-and-tumble locomotion to improve food capture in flowing environments

#### Abstract

#### Analysis of Direction-Dependent Speed across Flow Conditions

To evaluate whether the speed of Rhagovelia is affected by the direction of its motion relative to water flow, we analyzed its speed distribution across different heading angles. The insect's movement directions were binned into 8 angular sectors spanning 45° each, covering the full 360° plane. The polar plots below show the mean speeds of the insect in each direction under three flow conditions: low, medium, and high. Despite increasing flow speeds, we observed no significant directional bias in the insect's movement speed across the heading bins. This suggests that the influence of flow-induced drag on the insect's active propulsion is minor compared to its intrinsic locomotion speed (typically 200 mm/s), and remains below the resolution of our recording protocol. These findings support the conclusion that the observed motion is predominantly driven by active control rather than passive drag effects.

#### Model comparison of run-length distributions

For every trajectory we fitted three light-tailed candidate laws by maximum likelihood method;

1. Log-normal,
2. Gamma, and
3. Weibull

The Akaike Information Criterion is

$$\mathrm{AIC} = -2 \ln \mathcal{L}_{\max} + 2k,$$

with $k = 2$ free parameters for each model and $\mathcal{L}_{\max}$ representing the maximized values of the likelihood function. The model with the smallest AIC is considered the best; the strength of support for the others is given by the *AIC difference*

$$\Delta_i = \mathrm{AIC}_i - \mathrm{AIC}_{\mathrm{best}}.$$

By convention:

- $\Delta < 2$: models are indistinguishable;
- $4 \leq \Delta \leq 7$: much less support;
- $\Delta > 10$: essentially no support (Anderson and Burnham (2002)).

Across the 15 trajectories the Weibull model is preferred in 11 cases, the Gamma in 4. In every experiment where Weibull is selected the next-best model has $\Delta\mathrm{AIC} > 10$, indicating decisive support. Log-normal fits are never competitive ($\Delta\mathrm{AIC} \geq 6$). These results indicate that run lengths possess a characteristic scale rather than a scale-free (Lévy) tail.

© The Author 2023. Published by Oxford University Press. All rights reserved. For permissions, please

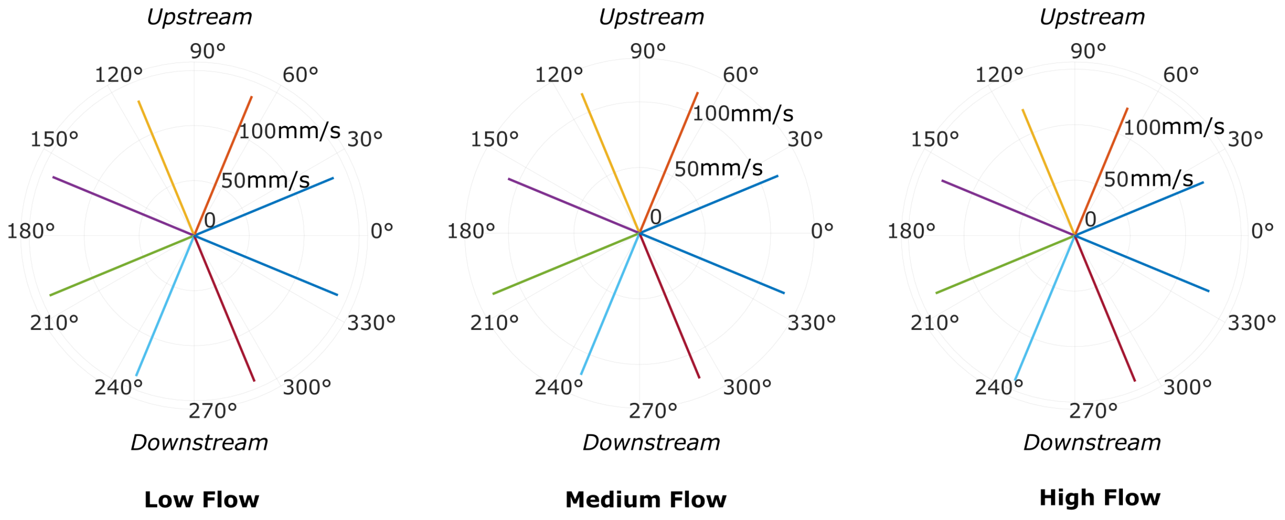


**Fig. 1. Direction-dependent speed analysis of *Rhagovelia* in flow.** Mean locomotion speed of the insect is plotted as a function of heading direction, divided into eight angular bins (each 45° wide), under three different background flow conditions: low, medium, and high. Across all flow regimes, no significant directional bias in speed is observed.

**Table 1.** AIC comparison for the five trajectories in each flow regime. BestAIC is the minimum AIC in that experiment; $\Delta AIC_{LN}$, $\Delta AIC_{G}$, and $\Delta AIC_{W}$ are the AIC differences for the log-normal, Gamma, and Weibull fits, respectively. The selected model (smallest AIC) is shown in the right-hand column.

| Flow | Exp | BestAIC | $\Delta AIC_{LN}$ | $\Delta AIC_{G}$ | $\Delta AIC_{W}$ | Selected |
|---|---|---|---|---|---|---|
| Low | 1 | 537.5 | 6.1 | 2.7 | 0.0 | Weibull |
| | 2 | 337.9 | 0.8 | 0.0 | 1.5 | Gamma |
| | 3 | 176.2 | 9.9 | 5.5 | 0.0 | Weibull |
| | 4 | 680.1 | 19.5 | 9.3 | 0.0 | Weibull |
| | 5 | 645.3 | 3.9 | 0.4 | 0.0 | Weibull |
| Medium | 1 | 978.5 | 17.4 | 6.2 | 0.0 | Weibull |
| | 2 | 310.0 | 1.6 | 0.0 | 1.5 | Gamma |
| | 3 | 1638.8 | 37.9 | 18.1 | 0.0 | Weibull |
| | 4 | 989.8 | 8.4 | 0.0 | 0.7 | Gamma |
| | 5 | 341.4 | 22.3 | 9.3 | 0.0 | Weibull |
| High | 1 | 2910.7 | 96.4 | 36.8 | 0.0 | Weibull |
| | 2 | 5609.8 | 233.5 | 140.8 | 0.0 | Weibull |
| | 3 | 3165.1 | 25.6 | 0.0 | 4.0 | Gamma |
| | 4 | 6461.0 | 94.3 | 14.3 | 0.0 | Weibull |
| | 5 | 2793.6 | 90.5 | 31.0 | 0.0 | Weibull |
